# GTC: a novel attempt to maintenance of huge genome collections compressed

**DOI:** 10.1101/131649

**Authors:** Agnieszka Danek, Sebastian Deorowicz

## Abstract

**Motivation:** *Results:* We present GTC, a novel compressed data structure for representation of huge collections of genetic variation data. GTC significantly outperforms existing solutions in terms of compression ratio and time of answering various types of queries. We show that the largest of publicly available database of about 60 thousand haplotypes at about 40 million SNPs can be stored in less than 4 Gbytes, while the queries related to variants are answered in a fraction of a second.

*Availability:* GTC can be downloaded from https://github.com/refresh-bio/GTC or http://sun.aei.polsl.pl/REFRESH/GTC.

*Contact:*

## 1 Introduction

In the last two decades the throughput of genome sequencers increased by a few orders of magnitude. At the same time the sequencing cost of a single human individual decreased from over 1 billion to about 1 thousand dollars. Stephens *et al.* (2015) predict that in 2025 the genomic data will be acquired at 1 zetta-bases/year, while about 2–40 exa-bytes/year of them should be deposited for a long term. From the other side, the prices of storage and transfer decrease moderately (Deorowicz and Grabowski, 2013), which means that keeping the data management costs under control becomes a real challenge.

Recently the tools for compression of sequenced reads have been benchmarked (Numanagić *et al.*, 2016). The ability of the examined utilities to shrink the data about ten times is remarkable. Nevertheless, much more is necessary. The obvious option is to resign from storage of raw data (in most experiments) and focus just on the results of variant calling, deposited usually in the Variant Call Format (VCF) files (Danecek *et al.*, 2011). The famous initiatives, like the 1000 Genomes Project (Sudmant *et al.*, 2015) or the 100,000 Genomes Project (The 100,000 Genomes Project, 2017), deliver VCF files for thousands of samples. Moreover, the scale of current projects, like of the Haplotype Reference Consortium (HRC) (McCarthy *et al.*, 2016) or the Exome Aggregation Consortium (ExAC) (Lek *et al.*, 2016), is by an order of magnitude larger. For example, the VCF files of the HRC consist of 64,976 haplotypes at about 39.2 million SNPs and occupy 4.3 TB. It is also clear that much larger databases will be formed in the near future.

VCF files contain a list of variants in a collection of genomes as well as evidence of presence of a reference/non-reference allele at each specific variant position in each genome. As they are intensively searched, the applied compression scheme should support fast queries of various types. The indexed and gzipped VCF files can be effectively asked using VCFtools (Danecek *et al.*, 2011) or BCFtools when the query is about a single variant or a range of them. Unfortunately, retrieving a sample data means time-consuming decompression and processing of a whole file.

Recently introduced Genotype Query Tools (GQT) (Layer *et al.*, 2016) made use of some specialized compression algorithm for VCF files. GQT was designed to compare sample genotypes among many variant loci, but did not allow to retrieve the specified variant as well as sample data. Shortly after that, Li proposed BGT (Li, 2015) based on the positional Burrows– Wheeler transform (Durbin, 2014). It offered more than 10-fold better compression than GQT and supported queries about genotypes as well as variants. Moreover, it allowed to restrict the range of samples according to some metadata conditions. The SeqArray library (Zheng *et al.*, 2017) for the R programming language is yet another solution to effectively compress and browse VCF files. The applied compression is based on the LZMA algorithm (Salomon and Motta, 2010).

In the present article we introduce GTC (GenoType Compressor) intended to store genetic variation data from VCF files in a highly compressed form. Our data structure, as confirmed by experiments, is much faster (often by an order of magnitude) in various types of queries. It is also significantly more compact than the competitors. These features allow the users to maintain huge collections of genotypes even at typical laptops with high speed of accession and could change the way the huge collections of VCF files are processed.

## 2 Methods

### 2.1 General idea and definitions

GTC is a new tool for compressed representation of genotypes supporting fast queries of various types. GTC compresses a collection of genotypes in a VCF/BCF format and allows for queries about genotypes:

- at specified range of variant sites positions (e.g., a single variant site), referenced to as variant query,
- of specified samples (e.g., a single sample), referenced to as sample query,
- combination of the above.

The ploidy of individuals determines the number of haplotypes that make up a single genotype. For diploid organisms, a genotype of an individual is defined by two separate haplotypes.

For precise description of the proposed algorithm let us introduce some terms. As an input we have a VCF/BCF file that describes *H* haplotypes at *V* single allele variants (sites). For any bit vector *X*, *X*[*i*] it the *i* th bit of this vector, *|X|* denotes its length.

### 2.2 Sketch of the algorithm

At the beginning, GTC divides the variants into blocks of some number of consecutive entries *b*_size_ (according to the preliminary experiments we picked the value *b*_size_ = 3584; a justification for such a choice can be found in the Supplementary Material) and processes each block separately (Fig. 1). The bit vectors representing presence/absence of reference alleles in all haplotypes are formed. In fact two bit vectors, referred to as variant bit vectors, are necessary to describe each variant, as four possibilities must be covered: (*i*) a reference allele, (*ii*) non-reference allele, (*iii*) the other non-reference allele, or (*iv*) unknown allele.

The haplotypes in each block are independently permuted to minimize the number of differences (i.e., a Hamming distance) between successive entries. As determination of the best permutation of haplotypes is equivalent to solving the Travelling Salesperson Problem (TSP) (Johnson and McGeoch, 1997) it is practically impossible to find the optimal solution in a reasonable time. Thus, the Nearest Neighbor heuristic (Johnson and McGeoch, 1997) is picked to quickly calculate a reasonably good solution. There are better algorithms for this task (in terms of minimizing the total number of differences between neighbor haplotypes). Unfortunately, they are too slow to be applied here. The description of the found permutation must be stored for each block to allow retrieval of the original data.

The permuted haplotypes are compressed (still within blocks) using a hybrid of specialized techniques inspired by Ziv-Lempel compression algorithm, run length encoding, and Huffman coding (Salomon and Motta, 2010). The variant bit vectors are processed one by one. In the current variant bit vector we look for the longest runs of 0s or 1s, as well as longest matches (same sequences of bits) in the already processed vectors. As the result we obtain a description of the current variant using the previous variants, which is much shorter than the original variant bit vector. The description is finally Huffman encoded to save even more space. These stages are described in detail in the following subsections, while Fig. 1 overviews a construction of GTC archive for an example VCF file (a more detailed overview can be found in the Supplementary Material).

### 2.3 Preprocessing the input VCF file

#### Managing the input VCF file

Unphased genotypes in the input BCF/VCF are arbitrary phased, while each of multi allele variant sites (described in a single line of VCF) is broken into multiple single allele sites, each described in a separate line, as in BGT VCF preprocessing (Li, 2015). Thus, there are four possible allele values for each haploid genotype: ‘0’ for the reference allele, ‘1’ for the non-reference allele, ‘2’ for the other non-reference allele (stored in a different line of the VCF file), ‘.’ for unknown (Fig. 1a).

#### Extraction of metadata

The altered description of *V* variant sites is stored in a site-only BCF file (the HTSlib (Li *et al.*, 2009) library is used for this task) with *_row* variable indicating site id added in the INFO field. List of names of samples is stored in a separate text file (Fig. 1b).

#### Initial encoding of genotypes

Each haploid genotype at each site is represented as a dibit (00 for the reference allele, 01 for the non-reference allele, 11 for the other non-reference allele, 10 for unknown (Fig. 1c)). Each of *V* site annotations is represented by two variant bit vectors of size *H*: one for lower and one for higher bits of the dibits representing successive haploid genotypes. Together there are 2*V* variant bit vectors (at this point information about all genotypes takes 2*HV* bits in total). The vectors are identified by ids ranging from 0 to 2*V -*1. The vectors with even ids correspond to the lower bits of the dibits representing haploid genotype sites, while vectors with odd ids correspond to the higher bits. In the next stages, described in details below, the variant bit vectors are compressed and indexed, making it possible to randomly access an arbitrary vector.

#### Forming blocks of genotype data

The variant bit vectors representing genotype data are divided into blocks. A single block contains genotype data of *b*_size_ = 3584 consecutive variant sites (value chosen experimentally) that is 2*b*_size_ = 7168 consecutive variant bit vectors. Thus, there are ⌈*V /b*_size_⌈ blocks (last may contain data about less than *b*_size_ variants). The blocks are processed independently to each other, in parallel (if possible).

### 2.4 Processing a single block of genotype data

#### Permutation of haplotypes

The haplotypes in a block are permuted to minimize the number of differences (i.e., a Hamming distance) between neighboring haplotypes. The Nearest Neighbor heuristic (Johnson and McGeoch, 1997) is used to calculate a reasonably good solution in an acceptable time. The permutation of the haplotypes (their order after permutation) in the *i*-th block is stored in *P*^*i*^ array.

#### Categorizing variant bit vectors

The variant bit vectors (representing genotype annotations) are processed one by one, in a sequential manner. They are packed into byte vectors (8 bits in a byte). A byte is then the smallest processing unit in the compression scheme. In the initial phase, each byte vector is categorized either as an empty vector (all bytes unset), a copy of a previously processed vector, or a unique vector, not equal to any of the previously processed vectors. The classification is done with help of a dictionary structure *HT*^vec^, namely hash table with linear probing (Knuth, 1998). The hash value is computed for each processed vector and *HT*^vec^ stores ids of all previously processed unique vectors in the block (notice: first non empty byte vector is a unique vector). Four bit vectors, each of Size 3584 bits, are formed for the *i-*th block: 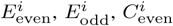 and 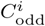 The *k*th set bit in 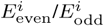%^%^ indicates that the *k*th even/odd (respectively) variant byte vector in the block is empty. The *k*th set bit%^%^ in 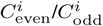 indicates that the *k*th even/odd (respectively) variant byte vector in the block is a copy of one of the previous vectors. The 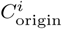 array maps subsequent copied vectors with their equivalents in the set of unique variant bit vectors (keeping id of unique vector, *uid*).

#### Processing unique variant bit vectors

Every unique variant bit vector, represented by byte vector (of size ⎡*H*/8⎤ bytes, padded with zeros), is transformed into a sequence of tuples, which represents literals, runs of zeros or ones, or matches to a previously encoded vector. The encoding is done byte by byte, from left to right, starting at position *j* = 0. At each analyzed position we first look for the length of the longest substring of zeros (bytes equal to zero) or ones (bytes equal to 255). If it is of length *r*_min_ or more (*r*_min_ = 2 by default), we encode the sequence as a run of zeros (or ones). Otherwise, we look for the longest possible match to one of the previously encoded unique vector (identical byte substring). To increase the chance of a long match, we search for common haplotypes, that is for the longest match starting at *j*th position in any previously processed byte vector. The matches to arbitrary parts of other vectors are accidental and thus unlikely to be long. Moreover, with the position restriction, the matches need fewer bits to encode (no need to store their positions). To make the search faster, the already processed data from the vectors are indexed using hash tables *HT*. Each *HT* ’s key consists of a sequence extracted from a vector and its position. The value is the unique vector id (*uid*). *HT* stores keys with sequences of length *h* = 5. The minimum match length is equal to *h* (and it is impossible to find shorter matches using *HT*). The matching byte substring (or its parts) in a previously encoded vector can be already encoded as a match. However, we restrict a *match depth* that is a number of allowed vectors describing each byte in a match. By default, the maximum allowed match depth (*d*_max_) is 100. A subsequent match to the same vector is encoded with fewer bits, as there is no need to store the id of the previous vector. If, with the match depth restriction, no sufficiently long match can be found, the current byte is encoded as a literal. The runs of literals (between 20 and 252 literals by default) are merged and encoded as a separate tuple.

The type of a tuple is indicated by its first field, a flag *f*_type_. Overall, there are six possible tuple types at the current position *j*:

- 〈*f*_zero_run_*, num*_0_〉—a run of *num*_0_ zero bytes, the position *j* is updated to *j* + *num*_0_,
- 〈*f*_one_run_*, num*_1_〉—a run of *num*_1_ one bytes (all bits set), the position *j* is updated to *j* + *num*_1_,
- 〈*f*_match_*, uid, len*〉—a match of length *len* to a vector with id *uid*, the position *j* is updated to *j* + *len*,
- 〈*f*_match_same_*, len*〉—a match of length *len* to a vector with the same id as the previous match, the position *j* is updated to *j* + *len*,
- 〈*f*_literal_*, bv* 〉—a literal, where *bv* is the value of the byte, the position *j* is increased by 1.
- 〈*f*_literal_run_, *n*, *bv_1_*, *bv_2_*, …, *bv_n_* 〉—a run of *n* literals, where *bv_1_*, *bv_2_*, …, *bv_n_* are the values of the consecutive bytes, the position *j* is increased by *n*.

### 2.5 Merging blocks

The ⌈*V /b*_size_⌉ processed blocks of genotype data are gathered and merged in the order of their appearance in the input VCF file (each *i*th block is added, for *i* = 1 to *i* = ⌈*V /b*_size_⌉).

The bit vectors 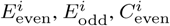, and 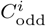 are merged into four global bit vectors of size *V*: *E*_even_, *E*_odd_, *C*_even_, and *C*_odd_. The vectors are kept in a succinct form.

The *C*_origin_ array gathers all 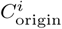%^%^ arrays adjusting stored *uid*s (number of unique vectors in all previous blocks is added to each *uid* from the current block). For subsequent copied vectors it refers to *uid*s of%^%^ the original unique vectors out of all unique vectors. Every *uid* in *C*_origin_%^%^ is then delta coded (difference between *uid* of the next unique vector and%^%^ original *uid* is calculated) with the minimum necessary number of bits.

The encoded unique variant bit vectors are stored in a byte array *U*. The flags, literals (*bv*), lengths of matches (*len*) and lengths of runs of zeros (*num*_0_) and ones (*num*_1_) are compressed (separately) with an entropy coder (we use Huffman coder) in the context of total number of set bits in the current variant bit vector (8 separate groups by default). For each literal run, the number of bits it occupies is kept. For matches, the ids of matched unique vectors (*uid*s) are delta coded with the minimum necessary number of bits.

The starting positions of all unique vectors in *U* are stored in a byte array *U*_pos_, where every 1025th unique vector is stored using 4 bytes, while for the 1024 subsequent vectors only the difference (between the current vector position and the nearest previous position encoded with 4 bytes) is stored, using *d* bits, where *d* is the minimum number of bits necessary to store any encoded position difference.

Finally, permutations of all blocks, *P*^*i*^, are merged into single array of permutations, *P*. It is stored with minimum necessary number of bits. The id of the variant bit vector to decompress is enough to find the right permutation in *P*.

### 2.6 Design of the compressed data structure

The main components of the GTC data structure are as follows:

- *E*_even_ and *E*_odd_: two bit vectors, each of size *V*, indicating if subsequent variant bit vectors (out of all vectors for lower or higher bits of corresponding haplotypes, respectively) are zero-only vectors,
- *C*_even_ and *C*_odd_: two bit vectors, each of size *V*, indicating if subsequent variant bit vectors (out of all vectors for lower or higher bits of corresponding haplotypes, respectively) are copies of other vectors,
- *C*_origin_: ids of the original vectors (out of all unique vectors) for successive vectors being a copy; delta coding and minimum necessary number of bits are used to store each id.
- *U*: byte array storing unique variant bit vectors, encoded into tuples and compressed (as described above),
- *U*_pos_: byte array with positions of the subsequent unique variant bit vectors in the *U* structure; full position is stored for every 1025th vector, the position of the remaining vectors are delta coded.
- *P*: byte array storing permutations of subsequent blocks; a single permutation is a sequence of ids of haplotypes reflecting the order in which they appear in the block. Each id is stored with minimum necessary number of bits (exactly: ⌈log2 *H*⌉ bits).

The *E*_even_, *E*_odd_, *C*_even_, and *C*_odd_ bit vectors are represented by the compressed structure (Raman *et al.*, 2007; Navarroand and Providel, 2012) implemented in the SDSL (Gog *et al.*, 2014) library.

### 2.7 Queries

By default, the entire collection is decompressed into VCF/BCF file. It is possible to restrict the query by applying additional conditions. Only variants and samples meeting all specified conditions are decompressed. The following restrictions are possible:

- range condition—it specifies the chromosome and a range of positions within the chromosome,
- sample condition—it specifies sample or samples,
- alternate allele frequency/count condition—it specifies minimum/maximum count/frequency of alternate allele among selected samples for each variant site,
- variant count condition—it specifies the maximum number of variant sites to decompress.

Two decompression algorithms are possible depending on the chosen query parameters. For most queries a variant-oriented approach is used. A sample-oriented approach is applied in queries about relatively small, i.e., up to 2000, number of samples, but only if decompressed range of variant sites span multiple blocks.

#### Variant-oriented

In this algorithm two vectors representing a variant site are fully decompressed. Their ids are known thanks to *_row* variable in the BCF with variant sites description. Initially, the *E*_even_, *E*_odd_, *C*_even_, and *C*_odd_ bit vectors are used to define a category of the variant bit vector. Decompression of an empty vector is straightforward. For a copied vector, the rank operations on *E*_even_, *E*_odd_, *C*_even_, and *C*_odd_ bit vectors are used to determine which copy is it, while the id of the original, unique vector is found using the *C*_origin_ array. The *U*_pos_ array is used to find position of the unique variant bit vector in the *U* array. The consecutive bytes of the variant bit vector are decompressed by decompressing and decoding all flags, literals, lengths of 0s and 1s runs, and matches. If a match is encountered, the variant bit vector containing the match is not fully decompressed. Instead, the complete decoding of all irrelevant tuples from the beginning of the vector up to the match position is skipped as far as possible. For example, stored bit length of a run of literal allows to skip the run without time-consuming literal decoding. The *P* array helps to find the original order of haplotypes. Only a subset of queried samples, if that is the case, is reported.

If a range of variant sites is queried, the decompression is speed up by keeping the track of an adequate number of previous, already decompressed unique variant bit vectors. Moreover, the permutation of haplotypes only needs to be read from the *P* array at the beginning of a block, not for each variant site separately.

#### Sample-oriented

In this approach all haplotypes of the queried sample are decompressed. For example, if sample is diploid, two haplotypes are decompressed. In case of more samples, more haplotypes are decompressed.

The decompression starts at the beginning of a block containing the first queried variant site. The *P* array is used at the start of each block to find the positions of bytes containing the haplotypes in the permuted variant vectors. Each variant vector is partly decoded once. The bytes containing information about the decompressed haplotypes are fully decoded and stored, the complete decoding of other bytes is skipped, if possible. As the previous decoded bytes are kept, if a match covering the decompressed haplotype is encountered (only a match within the same block is possible), the byte value can be read immediately. The decompressed bytes are used to retrieve values of queried haplotypes (a dibit, two single bits in two successive variant bit vectors).

## 3 Results

### 3.1 Data sets

To evaluate GTC and the competitors we picked two large collections of *H.sapiens* genomes: the 1000 GP Phase 3 (2,504 diploid genotypes) and the HRC (27,165 diploid genotypes). The 1000 Genomes Project data were downloaded from the project Web site (details in the Supplementary Material). The HRC collection used in our experiments was picked from the European Genom-phenom Archive (accession number: EGAS00000000029; details in the Supplementary Material) and is slightly smaller than the full data set (containing 32,488 samples; unfortunately unavailable for public use).

### 3.2 Compression ratios and times

The characteristics of the datasets and the compression capabilities of gzip, BGT, GQT, SeqArray, and GTC are summarized in Table 1. GTC archive appeared to be about two times more compact than BGT archive and tens times than gzipped VCF. The SeqArray archive size is between GTC and BGT for the smaller collection, but is noticeably larger than BGT for the larger one.

**Table 1.** Datasets used in the experiments.

| Dataset | No. genot. | No. var. [M] | Size [GB] |  |  |  |  |  |
| --- | --- | --- | --- | --- | --- | --- | --- | --- |
|  |  |  | VCF | VCF-gzip | GQT | BGT | SA | GTC |
| 1000GP-3 | 2,504 | 85.20 | 853 | 17.41 | 14.34 | 3.87 | 2.93 | **1.81** |
| HRC | 27,165 | 40.36 | 4,300 | 75.01 | 84.42 | 7.01 | 12.68 | **3.98** |
| HRC Chr11 | 27,165 | 1.94 | 210 | 3.42 | 4.11 | 0.33 | 0.62 | **0.18** |

The sizes of the compressed representations are also presented.

For a detailed examination of the compression ratios we reduced the HRC collection to 1000, 2000, etc. samples containing only Chromosome 11 variants. Figure 2a shows that the relative sizes of the BGT and GTC archives are similar in the examined range. A more careful examination shows that the compressed sizes of BGT and GTC grows slightly sub-linearly for growing number of genotypes. Moreover, the margin between BGT and GTC is almost constant for more than 5000 samples, which suggests that it would remain the same for even larger collections. SeqArray seems to scale poorer than BGT and GTC when the collection size grows.

**Fig. 1.**
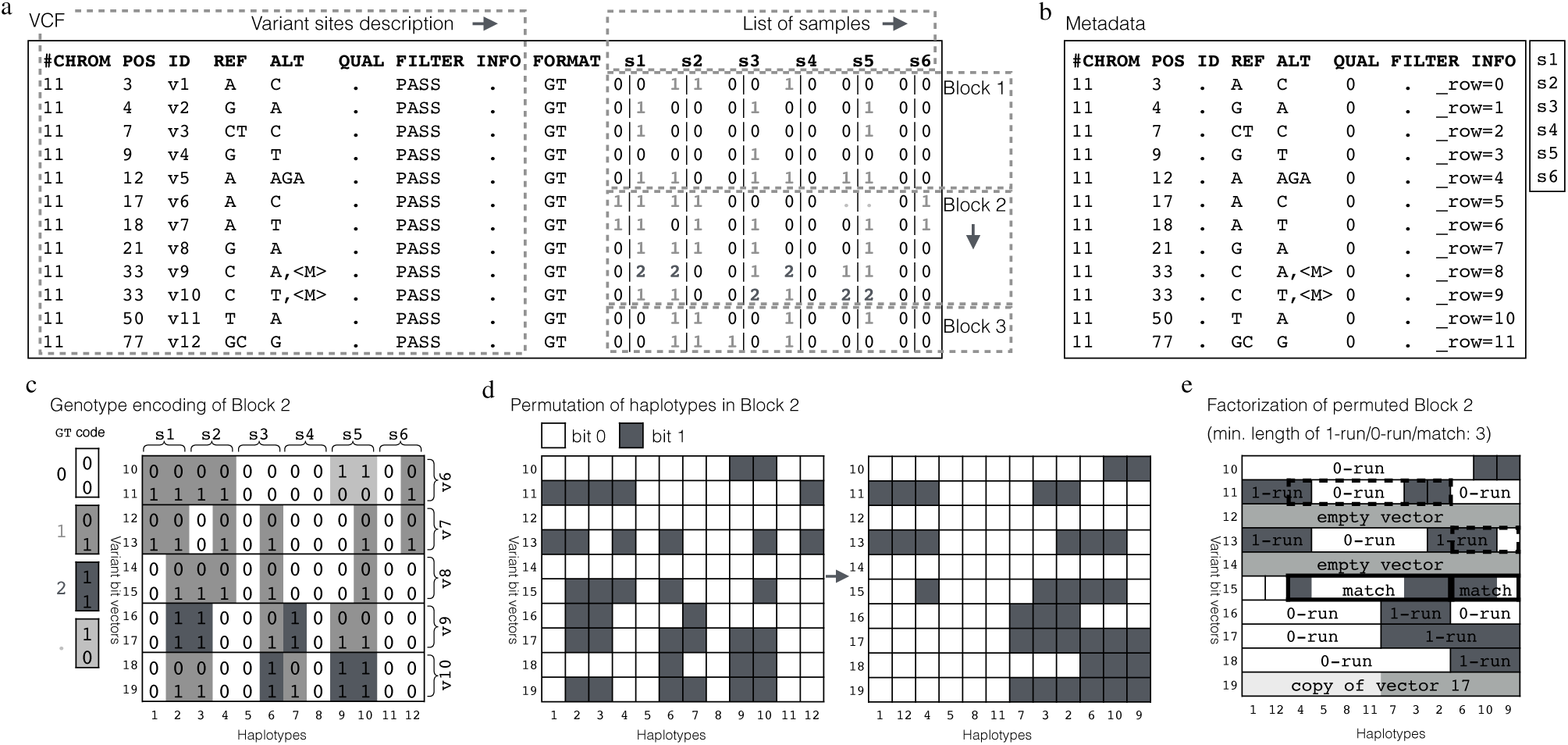
Construction of GTC archive. (**a**) The input VCF file with genotypes of 6 diploid samples at 12 variant sites. It is decomposed into metadata and blocks of genotypes. Each block (here: max. 5 variant sites) is processed separately. (**b**) Metadata: variant sites description (stored as a site-only BCF) and list of samples. (**c**) Bit vector representation of genotypes in Block 2. Each variant site is described by two variant bit vectors. The genotype of a haplotype, at each variant site, is described by two bits (located in two successive variant bit vectors). (**d**) Permutation of haplotypes in Block 2. Resulting order of haplotypes is stored. (**e**) Factorization of permuted Block 2. Empty and copied vectors are marked. All unique bit vectors are described as a sequence of longest possible 0-runs, 1-runs, matches to previous variant bit vectors in the block and literals. Note: The extended version of this example can be found in Supplemental Fig. 1.

**Fig. 2.**
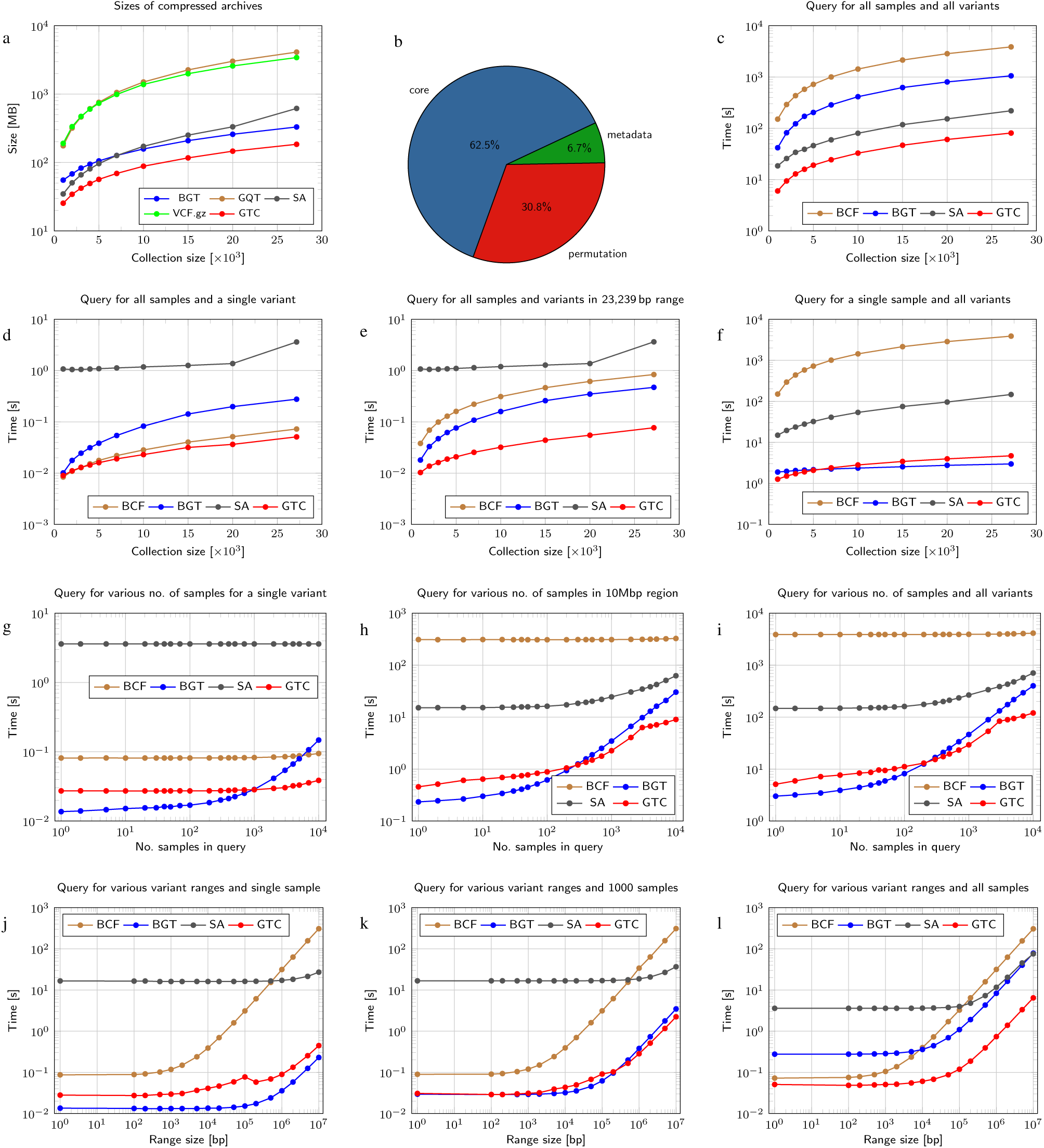
Comparison of compressed data structures for VCF (gzipped, VCF.gz) with BCFtools (BCF) used to access it, GQT, BGT, SeqArray (SA), and GTC. (**a**) Sizes of compressed sampled archives of HRC data. (**b**) Components of GTC archive for HRC data. (**c**) Decompression times for HRC Chromosome 11 data. (**d**) Single variant range query times for HRC Chromosome 11 data. (**e**) Variants in range of size 23,239 bp query times for HRC Chromosome 11 data. (**f**) Single sample query times for HRC Chromosome 11 data. (**g**) Single variant for selected samples query times for HRC Chromosome 11 data. (**h**) Variants in range of size 10 Mbp for selected samples query times for HRC Chromosome 11 data. (**i**) All variants for selected samples query times for HRC Chromosome 11 data. (**j**) Single sample for selected range of chromosome query times for HRC Chromosome 11 data. (**k**) 1000 samples for selected range of chromosome query times for HRC Chromosome 11 data. (**l**) All samples for selected range of chromosome query times for HRC Chromosome 11 data.

It is also interesting to see the sizes of components of GTC archive for the complete HRC data. As can be observed in Fig. 2b the majority of the archive (62.5%) is for the description of matches, 0-runs, etc. 30.8% is for the description of permutations in blocks. Finally, 6.5% is for the description of variants (position, reference and non-reference alleles, etc.) and 0.2% for list of sample names. For smaller block sizes the haplotypes can be better compressed, but the description of permutation consumes more space (Supplemental Fig. 8). In Supplemental Figures 2–7 we also show the influence of various parameters of GTC (like block size, minimal match length, etc.) on the compression ratio.

The compression times of BGT, SeqArray, and GTC are similar (slightly more than a day for the complete HRC collection at Intel Xeon-based workstation clocked at 2.3 GHz) as they are dominated by reading and parsing of the huge input gzipped VCF file.

### 3.3 Queries to the compressed data structure

The great compression would be, however, of little value without the ability of answering queries of various types.

Figures 2c-l show the times of answering various types of queries by GTC, BGT, SeqArray (SA), and BCFtools (BCF), for Chromosome 11 data containing various number of samples (various collection sizes, Fig. 2c-f) or for the whole collection (27,165 samples, Fig. 2g-l). The GTC decompression of the whole collection of variants is from 7 times (1000-sample subset) to 13 times (all samples) faster than of BGT and about 3 times faster than of SeqArray (Fig. 2c).

The GTC extraction of single variant or range of genome of size 23,329 bases (median size of human gene) is up to 6 times faster than of BGT (Fig. 2d-e). It is also worth to note that BCFtools are almost as fast as GTC for a single variant query.

A bit different situation can be observed in Fig. 2f, where the decompression times of single sample are presented. For collections not larger than 5,000 samples GTC is the fastest, but then, BGT takes the lead becoming 1.5 times faster for the complete collection. Nevertheless, both GTC and BGT answer the query in a few seconds. As expected BCFtools are clearly the slowest.

Figures 2g-l show the times of answering the queries about different ranges of genome and various number of samples for the complete Chromosome 11 collection (27,165 samples). The BGT extraction times are slightly better than of GTC only when a small number of samples (up to about 1 thousand samples for small variant ranges and up to few hundreds samples for large variant ranges) is queried. For more samples in a query, GTC takes the lead, while the relative performance of BGT decreases (i.e., for single variant and 10,000 samples extraction BCFtools are the second best method). When extracting all samples GTC is about ten times faster than BGT. Moreover, the advantage of GTC grows slightly when the range extends. It can be noted how the relative performance of BCTtools degrades when the range width exceeds a few thousands bases.

From the pure compression-ratio perspective it is worth to note that the average size of genotype information in GTC archive for a single sample is just about 146 KB. Even better result would be possible when we resign from fast queries support. To date the best pure compressor of VCF files is TGC (Deorowicz et al., 2013). Recently proposed GTRAC (Tatwawadi *et al.*, 2016) is a modified version of TGC, supporting some types of queries, e.g., for single (or range of) variants or single samples. Unfortunately, the output of GTRAC queries is just a binary file so no direct comparison with VCF/BCF-producing compressors (BGT, GTC, SeqArray) is possible. Moreover, we were unable to run GTRAC for the examined datasets. Nevertheless, we modified our compressor to produce similar output and support similar query types, and run both GTC and GTRAC for the 1000GP Phase 1 data containing 1092 genotypes and 39.7M variants. The compressed archives were of size 622 MB (GTC) and 1002 MB (GTRAC). The queries for single variants were solved in 7 ms (GTC) and 47 ms (GTRAC). The queries for samples were answered in similar times, about 2 s for Chromosome 11 data.

## 4 Conclusions

The widely used VCFtools and BCFtools offer relatively quick access to VCF, gzipped VCF, or BCF files when we ask about a specified variant or variant range. The situation, however, changes when we focus on sample queries, which are slow. Another drawback of using these tools is that the gzipped VCF or BCF files are quite large, which causes the problems with storage and transfer.

When the user is interested only in the genotype data from the VCF files much better solutions are possible, i.e., the state-of-the-art BGT and SeqArray. Nevertheless, the GTC, proposed in this article, has several advantages over them. The most significant one is a new compression algorithm designed with a care of fast queries support. The data structure is so small that it is possible to store it in the main memory of even commodity workstations or laptops giving the impressive query times. Even when run at powerful workstation, GTC often outperforms the competitors in query times by an order of magnitude. The scalability experiments suggest that much larger collections can be also maintained effectively.

